# Along-Tract Statistical Mapping of Microstructural Abnormalities in Bipolar Disorder: A Pilot Study

**DOI:** 10.1101/2023.03.07.531585

**Authors:** Leila Nabulsi, Bramsh Q. Chandio, Nikhil Dhinagar, Emily Laltoo, Genevieve McPhilemy, Fiona M. Martyn, Brian Hallahan, Colm McDonald, Paul M. Thompson, Dara M. Cannon

**Affiliations:** Imaging Genetics Center, Stevens Institute for Neuroimaging & Informatics, University of Southern California, Marina del Rey, CA, 90292 USA; Clinical Neuroimaging Lab, Galway Neuroscience Centre, College of Medicine, Nursing, and Health Sciences, University of Galway, Galway, Ireland

## Abstract

Investigating brain circuitry involved in bipolar disorder (BD) is key to discovering brain biomarkers for genetic and interventional studies of the disorder. Even so, prior research has not provided a fine-scale spatial mapping of brain microstructural differences in BD. In this pilot diffusion MRI dataset, we used BUndle ANalytics (BUAN), a recently developed analytic approach for tractography, to extract, map, and visualize the profile of microstructural abnormalities on a 3D model of fiber tracts in people with BD (N=38) and healthy controls (N=49), and investigate along-tract white matter (WM) microstructural differences between these groups. Using the BUAN pipeline, BD was associated with lower mean Fractional Anisotropy (FA) in fronto-limbic and interhemispheric pathways and higher mean FA in posterior bundles relative to controls. BUAN combines tractography and anatomical information to capture distinct along-tract effects on WM microstructure that may aid in classifying diseases based on anatomical differences.

## I. INTRODUCTION

Mental health disorders affect more than 1 billion people globally, causing 7% of all global burden of disease (as measured in disability-adjusted life years) and 19% of all years lived with disability. Serious mental illnesses including BD, directly impact nearly one in 25 adults in the United States [1]. Thus, reliably characterizing the neural substrates involved in BD, is essential to discovering brain biomarkers for genetic and interventional studies, for accurate treatment targeting, outcome prediction, and modeling of inter- and intra-individual disease variability. Bipolar disorder (BD) has been increasingly considered a *dysconnection syndrome* as a result of the complex interplay between gray and white matter (WM) components involving emotion-regulatory circuitries [2]. Contemporary neurobiological theories of emotion propose a constellation of fronto-limbic regions, and their connections and interactions with regions traditionally implicated in anxiety and fear circuitries as well as cognitive control. Corresponding models of BD suggest that dysfunction in these fronto-limbic circuits underpin the emotional and cognitive dysregulation that characterize the disorder. While prior research using structural MRI has employed voxel-based techniques such as VBM to characterize brain differences in white matter in BD [3] - despite the known limitations of VBM [4] - recent efforts employing diffusion MRI have not provided a fine-scale spatial mapping of microstructural differences but have instead relied on region-of-interest (ROI), tract-based spatial statistics (TBSS) [5], and more recently, network-based approaches [6].

Meta-analyses of studies investigating white matter connectivity in BD report abnormalities in major association pathways and predominantly limbic dysconnectivity as a signature of the illness. The first meta-analysis of DTI studies in BD reported lower fractional anisotropy (FA) in the superior longitudinal (arcuate) and inferior *fasciculi*, inferior fronto-occipital gyrus, and posterior thalamic radiation, as well as the limbic right anterior and subgenual cingulate cortex [7]. Nortje and colleagues (2013) meta-analyzed several voxel-based analyses (VBA) and implicated all major classes of tracts in the disorder [3]. Furthermore, meta-analysis of tract-based spatial statistics (TBSS) studies showed lower FA of right posterior tempo-parietal and two left cingulate clusters [3]. Lower FA has also been reported in the internal capsule, corpus callosum and uncinate *fasciculus* [8]. Lower global FA and white matter volumes in the callosum, posterior cingulum and prefrontal areas could explain compromised inter-hemispheric communication [9]. In contrast, higher FA has been reported in the *genu* of the corpus callosum [10] as well as bilateral inferior parietal/precuneus, and superior occipital white matter in BD [11]. A recent investigation of free water molecules in the extracellular space of neuronal tissue corroborated previous FA abnormalities seen in DTI studies [12]. The impaired white matter projections accommodate major fibers carrying impulses to and from the cortical areas that regulate emotional, cognitive, and behavioral aspects of BD; impaired white matter connectivity among posterior white matter tracts may represent a compensatory effect from disrupted frontal connectivity [13]. Graph-theory tools have been applied to whole-brain networks, with findings consistent with morphological disturbances in the corpus callosum, implicating inter-hemispheric structural dysconnectivity. Local structural and functional network alterations have been reported in prefrontal and limbic areas [6], [14], [15]. Network-based analyses (structural and functional) have also highlighted disturbances in broader circuit and network mechanisms that integrate cognitive control, affective and reward-systems of the brain [16]–[18]. An additional feature of white matter organization in BD is the presence of white matter hyperintensities, which may be caused by vascular ischemic processes [8], and may contribute to the reported white matter abnormalities. In summary, studies of white matter organization of BD suggest that abnormalities are not limited to anterior fronto-limbic pathways but are more widespread and may combine to mediate BD cognitive impairments and emotion regulation. These deficits may reflect a loss of coherence or differences in the number or density of white matter tracts or may suggest myelin/axonal disruptions.

The analysis of digital models of WM - or bundle recognition, extraction, or segmentation - has benefited from advances in tractography based on diffusion MRI. These advances make it easier to apply automated bundle segmentation methods in brain research to study pathways and gain insight into the alterations in WM microstructure that characterize diseased and healthy brains. In this pilot dataset, we used BUndle ANalytics (BUAN) [19] a recently developed analytic approach for tractography, to reconstruct refined anatomic maps along WM bundles from whole-brain tractograms in BD and healthy controls. BUAN has recently been applied to map and visualize the profile of microstructural abnormalities on 3D models of fiber tracts, yielding fine-scale maps of the effects of Parkinson’s disease [19], mild cognitive impairment (in ADNI3; [20]), and aging [21]. BUAN is also freely available in DIPY - https://dipy.org/documentation/1.5.0/interfaces/buan_flow/. Using BUAN, we aimed to investigate statistically significant along-tract WM microstructural differences between people with BD and controls. We hypothesized that people with BD would exhibit the greatest alterations in fronto-limbic and interhemispheric WM projections.

## II. METHODS

We analyzed cross-sectional 3D diffusion-weighted MRI data collected at 3T (HARDI; *b*=1200 s/mm^2^; 62 gradients; 1.8×1.8×1.9 mm voxels; FOV: 198×259×125 mm) from 38 individuals with BD (age: 38.8±13.2 y; 55% female) and 49 psychiatrically healthy controls (age: 44.5±12.2; 59% female). Scans were corrected for subject motion, including rotating the *b*-matrix, and corrected for eddy-current distortions (ExploreDTI v4.8.6) [22]. To account for crossing fibers within voxels, we used a deterministic (non-tensor) constrained spherical deconvolution (CSD) algorithm [23] (ExploreDTI). Several algorithms can be used to trace or reconstruct white matter trajectories based on the diffusion acquisition. While diffusion tensor imaging (DTI) allows for quantification of the diffusion coefficients – namely the displacement of water molecules in a preferential direction at every voxel in the brain – and has been mostly employed to date, it suffers from partial volume effects from smaller fiber orientations [24]. Thus, we used CSD, a more anatomically meaningful approach that better reflects the underlying microstructural organization of the fiber bundles. Diffusion eigenvector estimation was performed using the robust estimation of tensor by outlier rejection (RESTORE) approach [25]. Fiber tracking commenced in each voxel, continued with 1 mm step size, (2 mm)^^3^ seed point resolution, >30° angle curvature threshold, and 20–300 mm length, and terminated at a minimum fractional anisotropy (FA) of 0.2. FA, a measure of microstructural tissue organization, was calculated at each voxel. Each subject’s FA map was affinely registered to an FA template in MNI space using the registration framework provided in DIPY [26] with an implementation similar to ANTs [27]. Individual whole-brain tractograms underwent streamline-based linear registration (SLR) [28] to a bundle atlas template in MNI space [29]. 37 bundles were extracted from each subject’s whole-brain tractograms using the auto-calibrated version of *RecoBundles* and a standard WM tract atlas [19], [30]. We used the BUAN tractometry pipeline to locate significant microstructural group differences at specific locations along the length of the tract. A tract profile for each extracted bundle was generated, where FA values were mapped onto all the points of the streamlines in a bundle per subject (**Fig. 1**). Note that BUAN uses all the points on all the streamlines in the bundle profile creation as opposed to traditional tractometry methods that simplify each tract to a single mean streamline [31], [32]. Each point on every streamline is assigned a segment number based on its closest Euclidean distance to the model bundle’s centroid (coming from an atlas). Each subject’s bundle profile consisted of 100 segments along the length of the tract. Data per segment coming from two groups were analyzed using linear mixed models (LMMs). 100 LMM models were run per bundle type (one per segment along the length of the tract). In LMMs, FA was set as the response variable with the group, age, and sex as fixed effect terms and subject as a random effects term. The random effects models were used to properly account for the correlations between data points on the same streamline belonging to one segment for a specific subject. For this pilot study, we localized differences in specific segments of bundles based on the microstructural measure, FA, in BD relative to controls.

**Figure 1.**
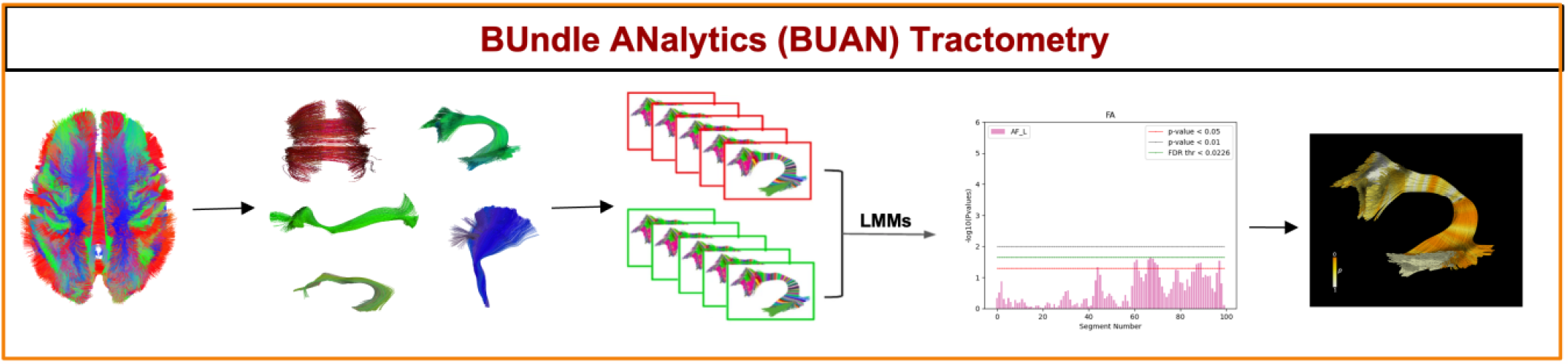
BUAN tractometry pipeline. White matter tracts are extracted from individual whole-brain tractograms; bundle profiles are created from the sets of extracted bundles from patients and controls. Bundle profiles are then analyzed using a linear mixed model, which yields a statistic of group difference at each segment along the tracts. These profiles can be displayed showing significant group deviations from control along the length of the tracts (see graph), and these *p*-values can also be textured back onto the 3D models of the tracts.

Multiple comparisons correction was performed controlling the False Discovery Rate (FDR) [33]. The correction was applied on each tract individually, accounting for 100 segments for each tract (\**p*_*FDR*_<0.05 to adjust for multiple comparisons while accounting for the non-independence of tract segments). FDR adjusted threshold is plotted for each tract in green in **Fig 1**.

## III. RESULTS

Our findings suggest that microstructural abnormalities in BD may extend beyond the white matter microstructural organization of fronto-limbic and inter-hemispheric connections that support emotion regulation in the brain to include posterior projections. Before multiple comparisons correction, compared to controls, the BD group exhibited lower FA within localized regions of the cingulum. Additionally, regions within the *fornix* and the corpus callosum (*forceps major* and *minor*) showed lower FA in the BD group, relative to controls. For significant segments, please see **Fig. 1, 2** and **3**.

**Figure 2.**
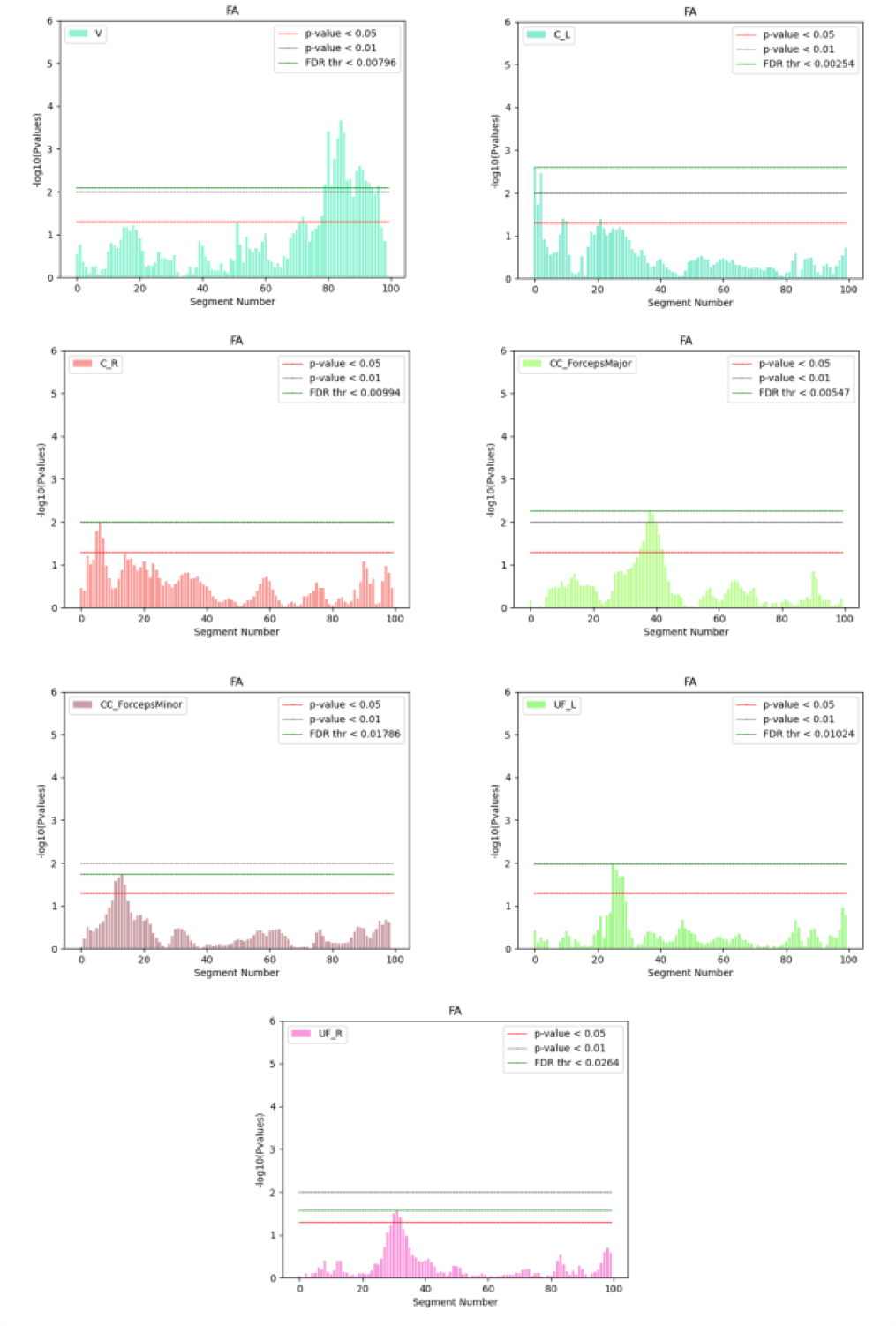
Localized along-tract microstructure (FA) alterations in BD. Compared to healthy controls, the group of BD participants exhibited altered FA in the following bundles (from left to right): V=Vermis; C=Cingulum; CC=Corpus Callosum; UF=Uncinate Fasciculus; ; L=left; R=right; FA=fractional anisotropy. In each plot, on the *x*-axis, segments along the length of the tract are shown and the *y*-axis shows the negative logarithm of the *p*-values. *P*-values that lie between or above two horizontal lines on the plot imply nominally significant group differences (uncorrected) at that location along the tract. The FDR adjusted threshold for each tract is plotted in green.

The arcuate (middle and inferior), longitudinal, and uncinate *fasciculi* also displayed lower FA in the patient group. Portions of the corpus callosum - and the right cingulum and uncinate bundles, which were also hypothesized to show effects - did not show significant group differences using FA in BD relative to controls. In the BD group, within a narrow subregion, higher FA was recorded in the brainstem, and specifically within the fronto-pontine, cortico-spino-thalamic, and medial lemniscus pathways, relative to controls (also illustrated in **Fig. 1** and **2**). Finally, in BD lower FA was detected in the vermis. Following multiple comparisons correction across all points on all tracts – a conservative correction – only localized regions within the vermis showed significant group differences. For 3D plots of significant bundles, see **Fig. 3**.

**Figure 3.**
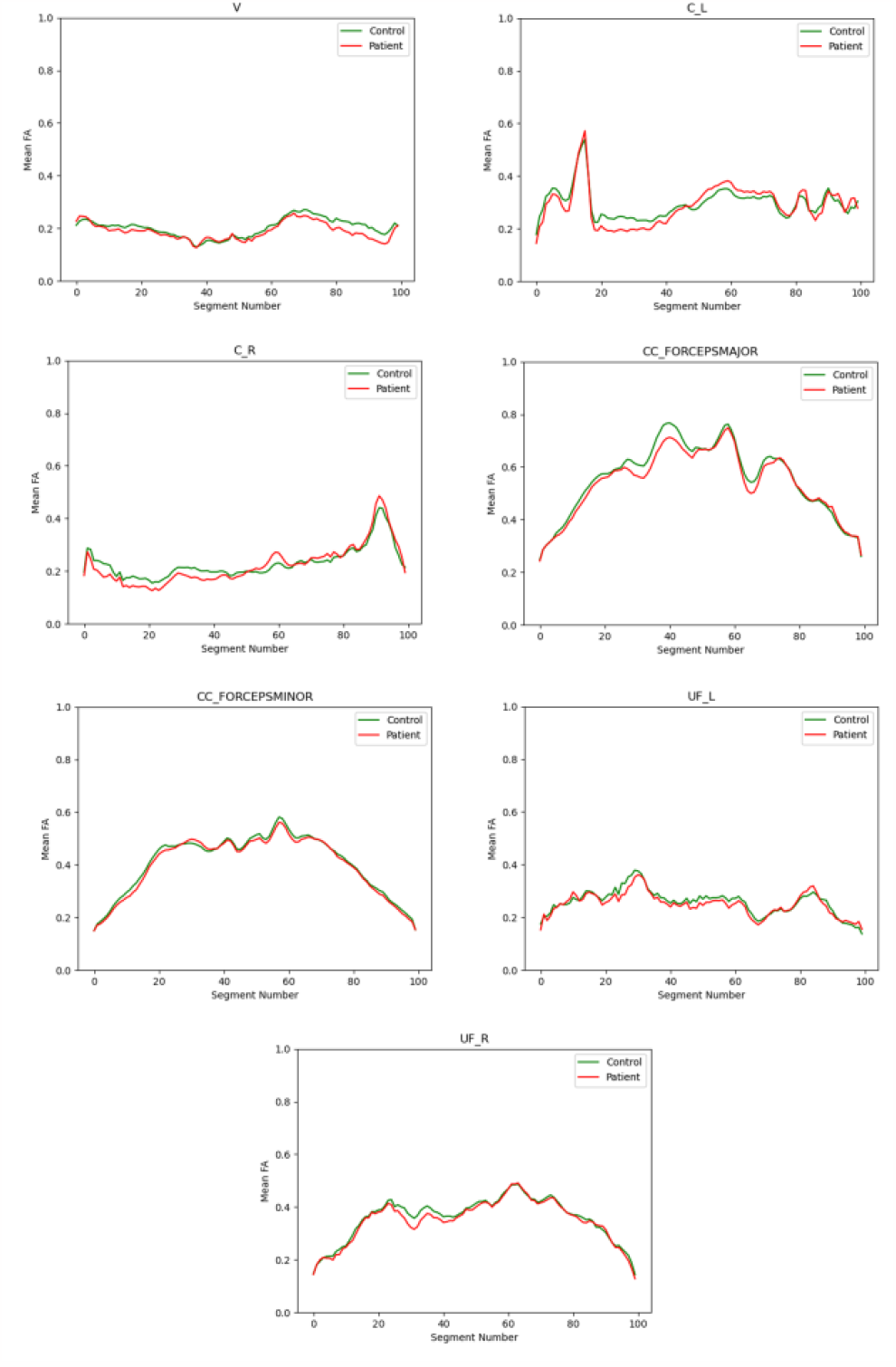
Group mean differences in fractional anisotropy in BD relative to controls along the trajectory of the bundles. Compared to healthy controls, participants with BD exhibited significantly higher FA in localized regions of the vermis, and lower FA in the cingulum, corpus callosum (*forceps major* and *minor*), and the uncinate fasciculus. In each plot, values of mean FA for controls are depicted in green, and in red for patients. For a key to the tract abbreviations, please see **Fig. 2**.

**Figure 4.**
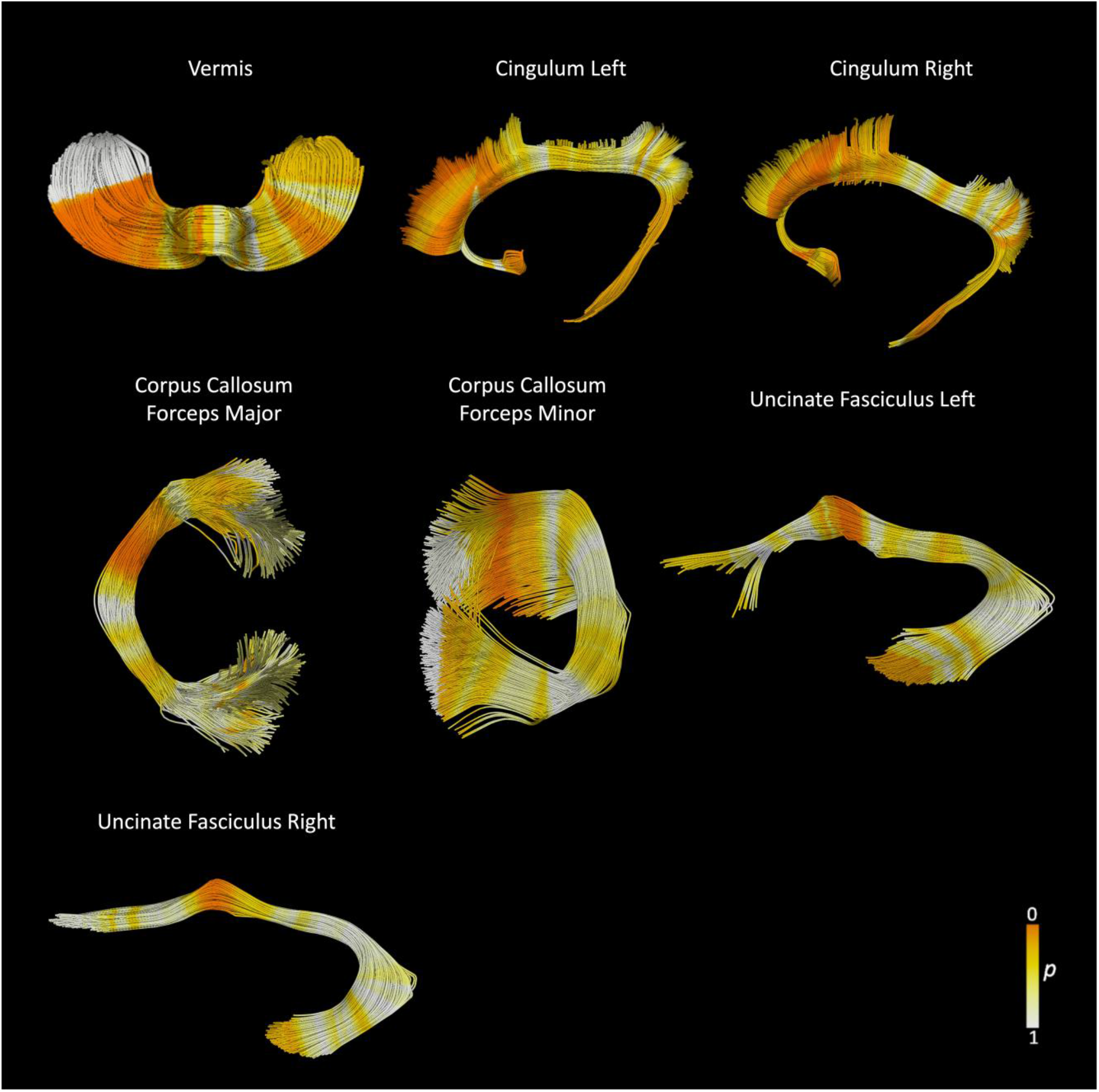
3D representation of group differences in WM bundles. Compared to healthy controls, the BD group exhibited microstructural differences in focal regions of 21 bundles; here, we present 7 bundles that are relevant to BD. Significant *p*-values (before FDR-correction) are depicted in *dark orange*. Following FDR-correction, group differences were recorded in localized regions of the vermis.

## IV. DISCUSSION

In this pilot study, we applied the BUAN tractometry pipeline to map, extract, and visualize the effects of BD on the white matter tracts of the brain. We found significant microstructural differences in cerebellar pathways in BD, relative to controls. Segments within fronto-limbic and interhemispheric projections also differed in the BD group compared to controls in terms of white matter microstructure. Research on the brain correlates of mental disorders has often focused on cortical regions and select subcortical structures, even though the human cerebellum contains 80% of the brain’s neurons. The cerebellum has been widely studied for its role in motor control, though through its extensive connections with association regions of the cortex the cerebellum is also integral to cognitive and affective processes. As cerebellar dysfunction is present in psychiatric disorders, it may contribute to the manifestation of affective and psychotic psychiatric symptoms [34]. Some anatomical studies report volumetric alterations in the cerebellum in BD [8]. Network-level analyses in BD suggest impaired microstructural organization in posterior-parietal regions, whereby the cerebellum was found to be less frequently involved in BD *rich club* membership [14]. Rich-club members are found in topologically central regions that are critical for global integration and coordination of higher cognitive processes within the brain’s networks [35]. Our finding is also consistent with network-level reports of functional dysconnectivity within the parieto-occipital and default mode network loops in BD [3], [7], [36]. Clinical variables describing symptom severity in BD (such as the number of episodes and duration of illness) have been associated with abnormal cerebellar connectivity; additionally, psychotic features have been linked to the cerebello-thalamo-prefrontal cortex network in BD [37], [38]. Despite this evidence, there are few reports of impaired white matter connectivity in BD involving the cerebellum. Future studies of white matter in BD should expand on existing cerebral-centric brain models of mental function by mapping cerebellar structure, as well as functional connectivity. Our regional findings implicate fronto-limbic and interhemispheric connections in BD. FA alterations, in anterior sections within the cingulum, are in regions where activations are detected in emotion-related fMR studies in BD [39], among other morphological and connectivity studies [16]. Additionally, alterations in the corpus callosum have been consistently found in structural, diffusion and functional MR studies of BD, particularly within the anterior horn (connecting bilateral prefrontal and limbic regions) [16]. The signal detected within the uncinate *fasciculus*, anterior to the *limen insula*, is also of interest to BD pathophysiology. It anatomically connects the orbitofrontal cortex to the anterior temporal lobes, participates in the amygdala-ventral prefrontal cortex system, and has been implicated in several psychiatric disorders. Traditionally considered as part of the limbic system, the uncinate may play a role in episodic memory, language and social emotional processing [40]. Recently, abnormalities in microstructural organization have been reported for the uncinate in the offspring of patients with BD; specifically, FA changes in the uncinate, over a follow-up period of 6 years, were found to predict the onset of BD illness [41]. While these findings may be of clinical interest, they should be considered exploratory, as they did not survive multiple comparisons correction (except for group differences in the cerebellum). To confirm the anatomical location of these microstructural alterations, this pilot study warrants replication in a larger sample. In our study, the FDR adjusted threshold was calculated per tract for 100 segments. However, proper adjustment for multiple comparisons may need to consider the autocorrelations in signals at adjacent segments along the length of the tracts. In future work, this will be addressed using functional data analysis [42] where the entire streamline will be treated as a function instead of testing segments independently [43]. This model may also enhance our statistical power and help validate our findings. Fractional anisotropy is the most consistently reported tensor metric and reflects the degree of anisotropic diffusion (or level of organization) within a fiber bundle [24], [44]. The presence of intact cell membranes and myelination may modulate anisotropy [45], thus differences in FA, may reflect neuroinflammation or changes in myelination [24], [46]. However, FA interpretation is controversial as it does not specifically associate with any single component of the underlying microstructure. Other tensor-derived metrics such as medial (MD), radial (RD) and axial (AD) diffusivity, measuring other directional aspects of the diffusion profile, have been employed as an alternative measure of white matter organization. Future studies will include these metrics in the context of BUAN tractometry to further characterize localized anatomical abnormalities in BD. Furthermore, the diffusion signal reflects several underlying contributions such as the degree of axonal myelination, extracellular differences in the medium leading to free water from inflammation and connectivity geometry (e.g., crossing fibers). While FA changes may reflect microstructural changes; these effects may be confounded by the above-mentioned factors. Here, CSD was used as it can reconstruct multiple diffusion directions within each voxel [47], [48]. Tractography-related strengths of this study include recursive calibration of the response function to better resolve volume effects within voxels [49]; rotation of the b-matrix during subject motion and eddy current correction steps to reduce errors in reconstruction in the orientation of the diffusion tensor [50]. While CSD approaches may propagate spurious reconstructed streamlines, the use of a stringent tracking angle, alongside FA thresholds (such as those employed here) should limit this possibility.

## V. CONCLUSION

Using an advanced along-tract analytic method, BUAN, we performed fine-scale spatial mapping of regional WM microstructure differences in BD, relative to controls. The BUAN tractometry pipeline successfully mapped and visualized the effects of BD on the white matter tracts of the brain. Significant microstructural group differences were observed in the vermis where FA decreases in the BD group, relative to controls. Before multiple comparisons correction, we also detected localized abnormalities involving fronto-limbic and interhemispheric pathways, corroborating focal findings in anterior and commissural fibers in BD. Additionally, white matter microstructure in cortico-thalamic projections may be also affected in BD. Intriguingly, significant group differences were found within posterior-cerebellar pathways in BD; these differences, may represent a compensatory mechanism to the dysconnectivity observed across fronto-limbic and inter-hemispheric projections when accounting for the microstructural organization of the bundle. The tracts implicated here connect regions that play important functional roles in the regulation of emotions, motivation, decision-making, and cognitive control [15], [18], which are impaired in BD. This pilot study, which requires replication, suggests that WM microstructure abnormalities in BD vary across an individual tract. This work highlights the complexity and functional specificity of fiber groups within complex large white matter bundles, and may guide future work characterizing WM in BD.

## ACKNOWLEDGMENT

Health Research Board (HRA-POR-324) and NIH grants RF1AG057892, R01MH116147, and R01MH129742.

